# Mathematical modelling of cortical neurogenesis reveals that the founder population does not necessarily scale with neuronal output

**DOI:** 10.1101/206045

**Authors:** Noemi Picco, Fernando García-Moreno, Thomas E. Woolley, Philip K. Maini, Zoltán Molnár

**Affiliations:** St John’s College Research Centre, St John’s College, St Giles Street, Oxford, OX1 3JP, UK; Wolfson Centre for Mathematical Biology, Mathematical Institute, University of Oxford, Woodstock Road, Oxford, OX1 6GG, UK; Department of Physiology, Anatomy and Genetics, University of Oxford, South Parks Road, Oxford, OX1 3QX, UK; Achucarro Basque Center for Neuroscience, Parque Científico UPV/EHU Edif. Sede, E-48940 Leioa, Spain; IKERBASQUE Foundation, María Díaz de Haro 3, 6th Floor, 48013 Bilbao, Spain; Cardiff School of Mathematics, Cardiff University, Senghennydd Road, Cardiff, CF24 4AG, UK

**Keywords:** Asymmetric division, cell cycle of cortical progenitors, cortical development, evolution, radial glial cells

## Abstract

The mammalian cerebral neocortex has a unique structure, composed of layers of different neuron types, interconnected in a stereotyped fashion. While the overall developmental program seems to be conserved, there are divergent developmental factors generating cortical diversity amongst mammalian species. In terms of cortical neuronal numbers some of the determining factors are the size of the founder population, the duration of cortical neurogenesis, the proportion of different progenitor types, and the fine-tuned balance between self-renewing and differentiative divisions. We develop a mathematical model of neurogenesis that, accounting for these factors, aims at explaining the high diversity in neuronal numbers found across mammalian species. By framing our hypotheses in rigorous mathematical terms, we are able to identify paths of neurogenesis that match experimentally observed patterns in mouse, macaque and human. Additionally, we use our model to identify key parameters that would particularly benefit from accurate experimental investigation. We find that the timing of a switch in favor of symmetric neurogenic divisions produces the highest variation in cortical neuronal numbers. Surprisingly, in our model the increase in cortical neuronal numbers does not necessarily reflect a larger size of founder population, a prediction that identified a specific need for experimental quantifications.

## Introduction

The mammalian cerebral neocortex has a unique structure, composed of layers of different neuron types, interconnected in a stereotyped fashion (Brodmann 1908; Cajal 1909; Rakic 2008; Amunts and Zilles 2015). The cortex is linked to numerous cognitive functions such as language, voluntary movement, and episodic memory. Understanding its development and evolution is key to shedding light on the divergent mechanisms that give rise to inter-species differences or diseases, such as microcephaly. The most striking differences across species consist of final neuronal output (i.e. the number of neurons produced by neurogenesis) and surface area (Krubitzer and Kaas 2005; Dehay and Kennedy, 2007), while the variation in radial thickness of the cortex is less pronounced (Rakic 2009, see Table 1). The 1000-fold variation in cortical surface area between mouse and human results in a remarkable specialisation and diversification of cortical function, allowing faster processing of complex information (Kaas 2013). However, the early developmental program that gives rise to each of these cortices seems to be conserved (Bystron et al. 2008; Lui et al. 2011; Kwan et al. 2012). Among the many factors determining the development of a normal cortex in mammalian species are: the size of the founder population present at the beginning of cortical neurogenesis, the length of neurogenesis, the dynamics of cell cycle length, the proportion of different cortical progenitor types, and the fine-tuned balance between self-renewing and differentiative divisions (Rakic 1995; Florio and Huttner 2014; Imayoshi and Kageyama 2014; Dehay and Kennedy, 2007). Qualitative changes in the outcome of neurogenesis following variations in any of these factors can be intuitively determined. For example, many decades ago Rakic predicted that a prolongation of the time devoted to amplification of the founder population results in expanded neuronal number hence cortical surface (Rakic 1988). However, understanding how all of these factors work together to ensure the cortex is correctly formed, finding the key parameters that has the greatest impact on the neuronal numbers is less amenable to verbal reasoning and more suitable for rigorous mathematical modelling.

**Table 1.**
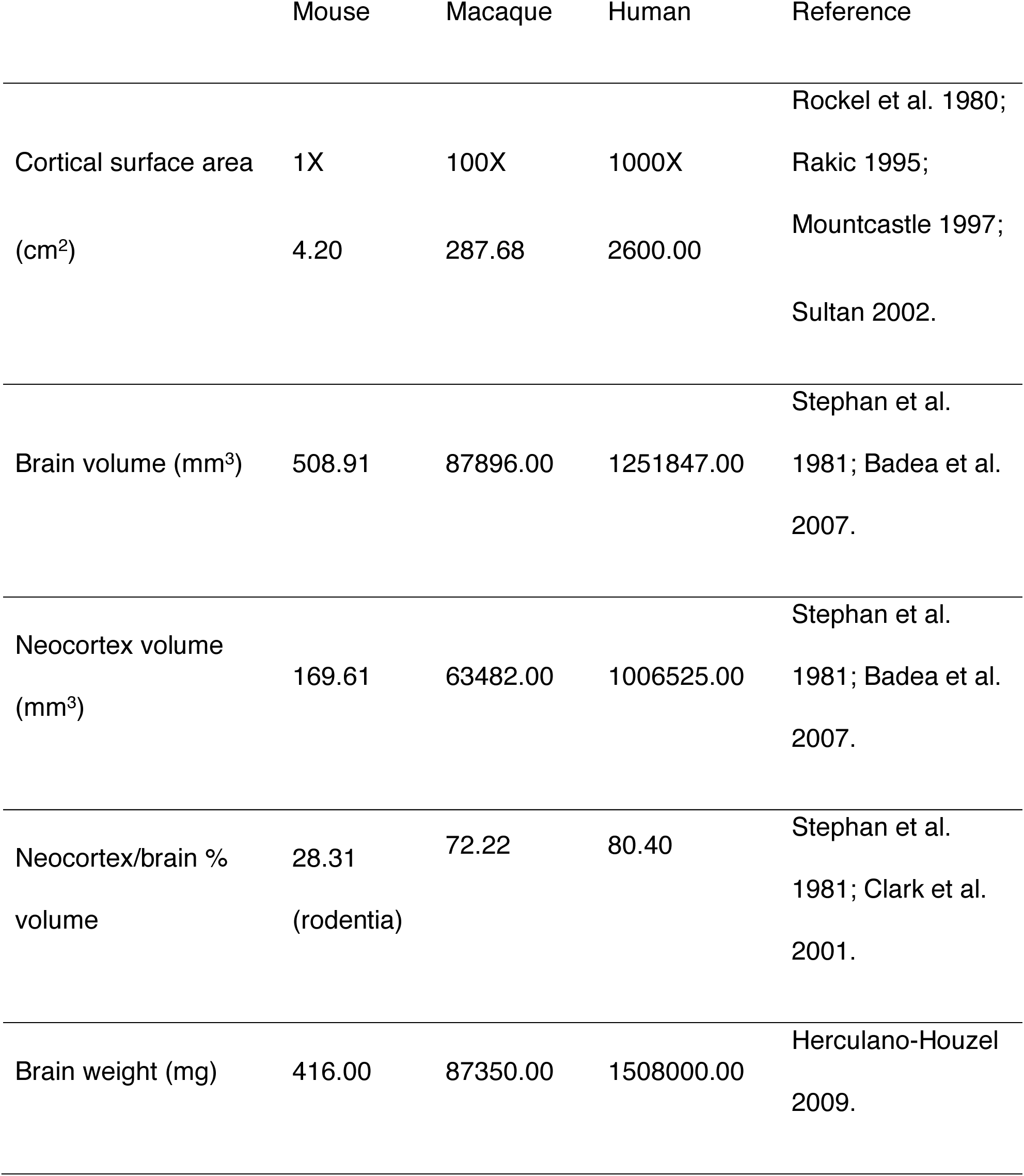
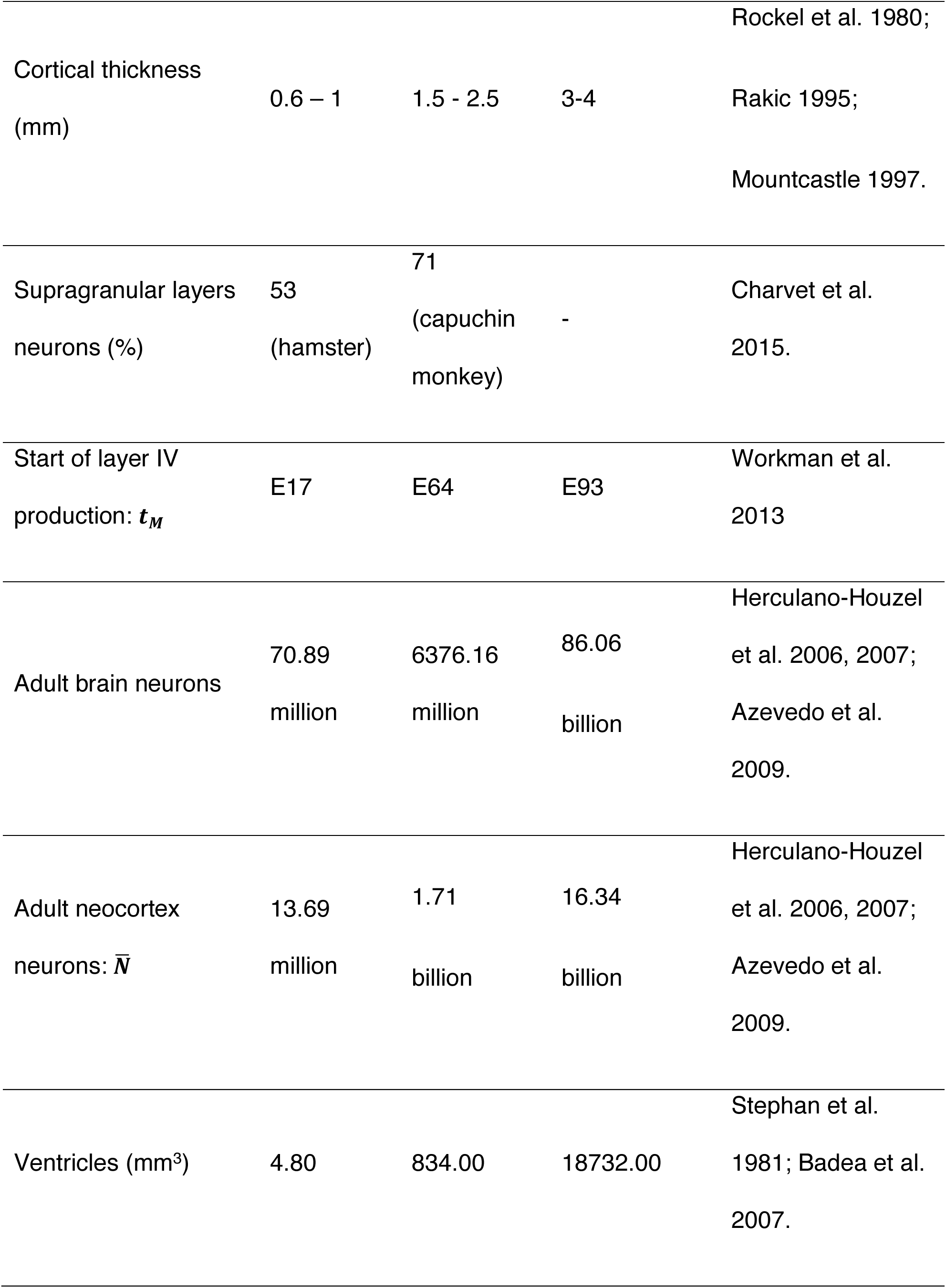
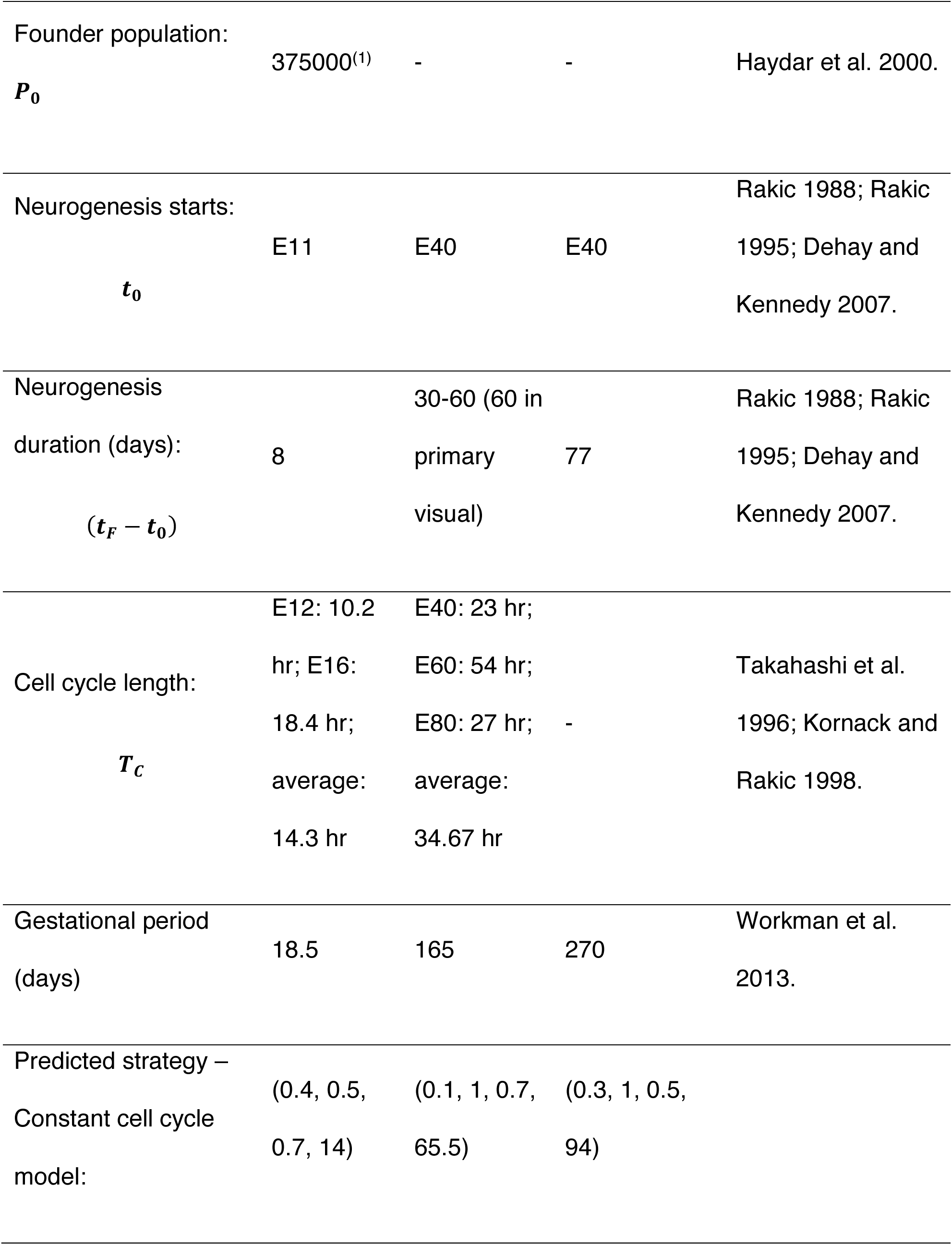
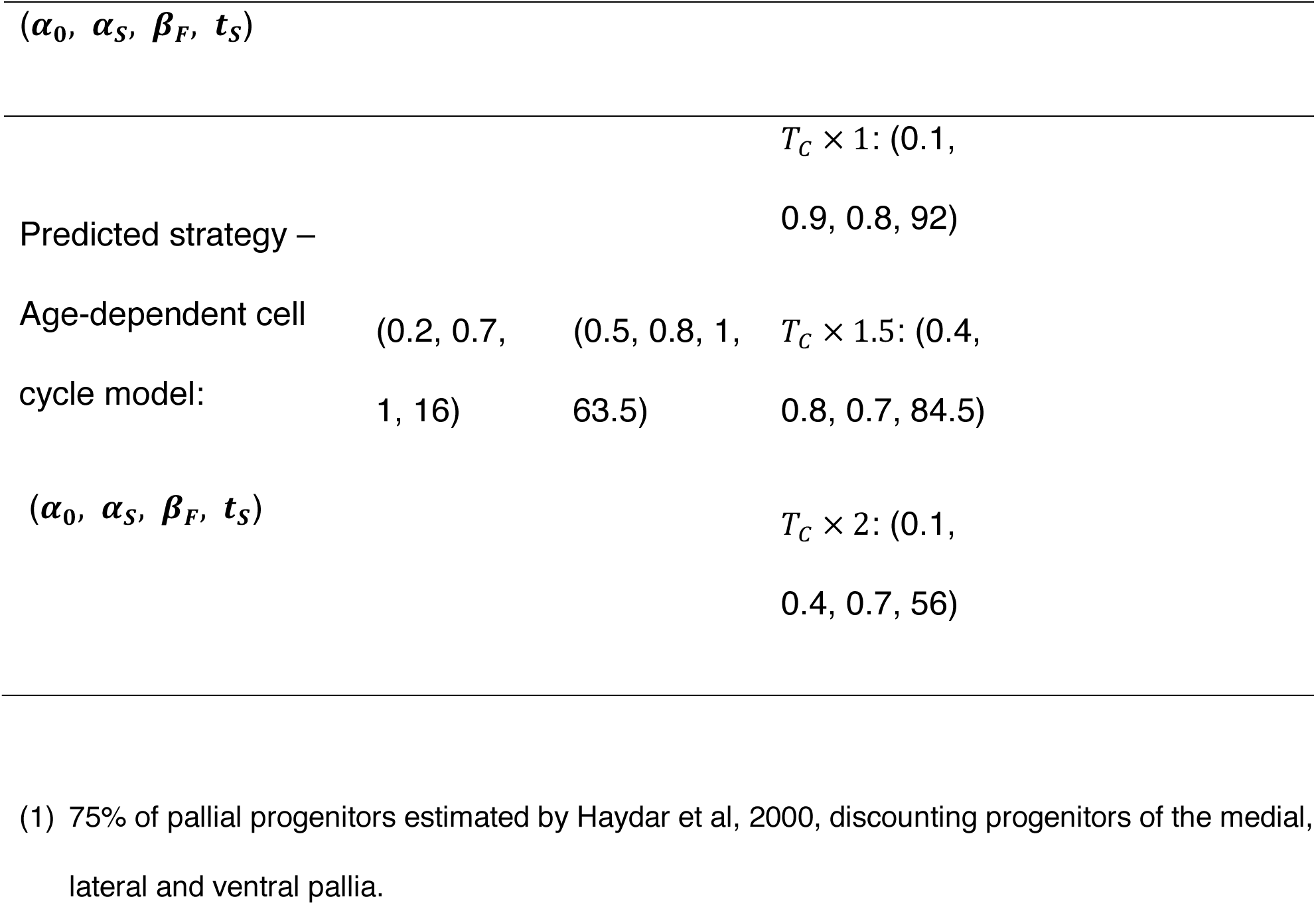
Quantities of interest for mouse macaque and human. E indicates embryonic day. (1) 75% of pallial progenitors estimated by Haydar et al, 2000, discounting progenitors of the medial, lateral and ventral pallia.

An impressive amount of comparative studies, quantifying and analysing the main differences in cortical neurogenesis of mammalian species, is available (Molnár et al. 2006; Cheung et al. 2007; Herculano-Houzel 2009; Martínez-Cerdeño et al. 2017. Some comparative quantitative data are beginning to be available on neuronal and glial cell numbers in various species (Herculano-Houzel 2012). However, these studies are limited to finding scaling rules that explain the variation in neuronal output by means of other observable characteristics, such as length of neurogenesis and brain volume, and do not investigate the underlying mechanisms (Mota and Herculano-Houzel 2015).

Recently, focus has turned to the identification of the progenitor types involved in cortical neurogenesis, in an attempt to exhaustively classify the cell types, from neural stem cells to post-mitotic neurons, and all stages of differentiation in between, in representative species such as mouse (Gal et al. 2006; Stancik et al. 2010), macaque (Wang et al. 2011; Betizeau et al. 2013), and human (Fietz et al. 2010; Hansen et al. 2010). It is possible to draw similarities in the process of cortical neurogenesis in mouse and human, such as the radial expansion of the cortex orchestrated by stage-specific dividing cells, resulting in the formation of the layers in an “inside-first, outside-last” fashion. However, the diversification of progenitor cell types, and non-apically constrained proliferation, are indicative of divergent processes adopted by different species in the course of evolution (Reillo et al. 2011; Pilz et al. 2013). The more we know about the processes involved in each species’ cortex development, the further we seem to be from finding a unifying theory that could explain the key variation observed in various mammals with cerebral cortex.

We propose a theoretical framework, based on a first-principle approach, to build a general, yet accurate, representation of a minimal set of processes and players involved in the cortical neurogenesis of the mammalian brain. Our model is minimal in the sense that we start from the simplest possible representation and introduce the time-dependency that is sufficient to capture the experimentally observed actual progenitor and neuronal numbers. Building on the current understanding of the processes involved, and framing hypotheses of unknown processes in a mathematically rigorous way, we can make testable predictions and/or expose where there is a lack of biological data necessary for a systematic testing of these hypotheses.

The final goal of this study is to integrate mathematical modelling and experimental observations to highlight both the communal processes in the early development of, and the divergent factors that can explain cortical diversity amongst, mammalian species.

## Materials and Methods

We define a mathematical model of cortical neurogenesis, describing the dynamics of proliferation and differentiation at the cell population level, during the development of a mammalian brain. For this model to be descriptive of any species considered, we classify cells into two cellular populations: progenitors (*P*) and neurons (*N*). The former includes all pre-mitotic neural progenitor cell types involved in neocortex development for a given species, e.g. neuroepithelial cells, apical and basal radial glial, intermediate progenitors (Rakic 1995; Huttner and Brand 1997; Noctor et al. 2004; Florio and Huttner 2014). The neuron population includes all post-mitotic and permanently differentiated cells. In our model we only consider neurons locally generated by progenitors situated between the ventricular and pial surfaces. Migration of glutamatergic neurons from outside sources (Barber and Pierani 2016; García-Moreno et al., 2018) or GABAergic neurons from subpallial ganglionic eminences (De Carlos et al. 1996; Parnavelas 2000; Anderson et al. 2001; Tamamaki et al. 2003) is not explicitly modelled, however it will be accounted for when integrating experimental quantifications into the model.

We consider three possible modes of progenitor cell division: self-amplifying division (SymP), generating two identical progenitors; asymmetric neurogenic division (AsymN), generating one progenitor and one neuron; symmetric neurogenic division (SymN), generating two neurons.

As we will see later, if these three types of division occurred with constant proportions, we would observe one of two possible outcomes: extinction or unlimited growth of the progenitor population. This type of model captures the eventual depletion of the progenitor population observed in adult cortical neurogenesis. Indeed, constant proportions of proliferation and differentiation were used to model similar dynamics in the adult hippocampus (Ziebell et al. 2014). It is known, instead, that these modes of proliferation are preferentially balanced at different stages of neocortical neurogenesis during development (Noctor et al. 2004; Götz and Huttner 2005), and this program seems to be conserved across species (Kornack 2000). Furthermore, the progenitor population is not depleted during embryonic neurogenesis, to allow for gliogenesis later in development (Mallamaci 2013; Kessaris et al. 2006).

We, therefore, define time-dependent probabilities describing the preferred mode of division in the progenitor population during neurogenesis: *α*(*t*) is the probability of AsymN at a given time *t*, *β*(*t*) is the probability of SymN; hence 1 − *α*(*t*) − *β*(*t*) is the probability of SymP. The modulation of these probability functions in time defines a strategy that can be represented as a trajectory in the three-dimensional space (AsymN, SymN, SymP) (Fig. 1).

**Figure 1.**
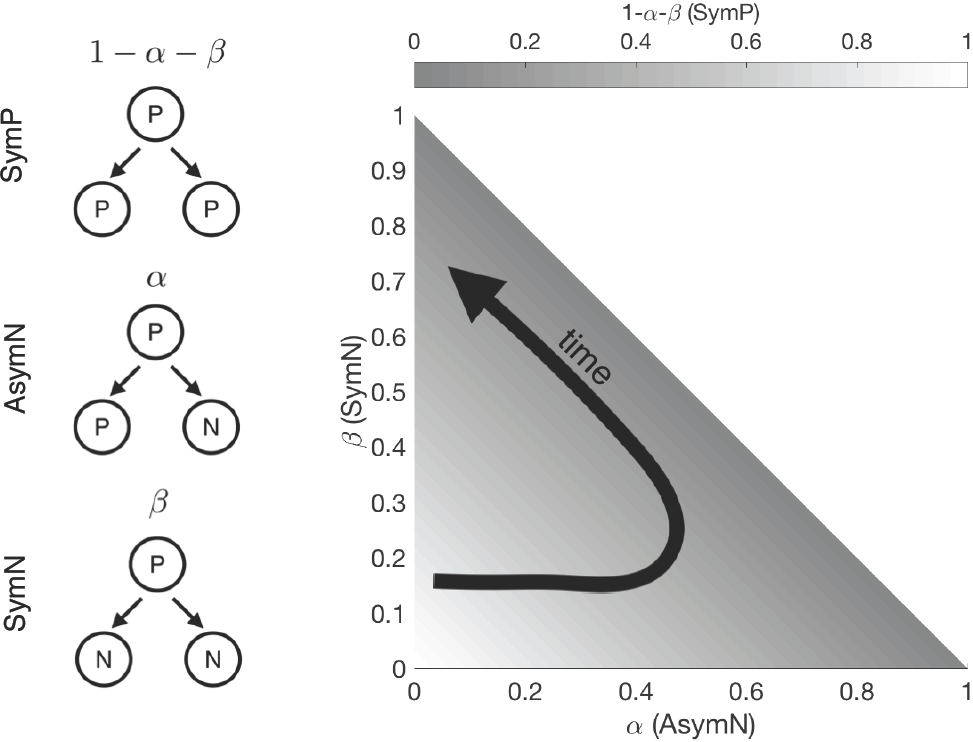
Schematic of the three types of cellular division (left) and corresponding strategy space (right). P = progenitor. N = neuron. The progenitor cell population navigates the strategy space by balancing probabilities of committing to each of the three division types: symmetric proliferative (SymP), asymmetric neurogenic (aSymN) and symmetric neurogenic (SymN). The trajectory indicated by the arrow is an example of time-dependent strategy, initially increasing the prevalence of AsymN, while reducing SymP, and finally favoring SymN.

The resulting system of Ordinary Differential Equations (ODEs), describing the evolution in time of the progenitor (*P*) and neuron (*N*) populations is:

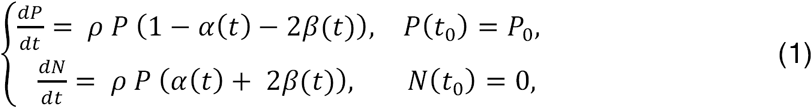

where *ρ* is the rate of division, and *P*_0_ is the founder population. We define *t_0_* as the time of onset of neurogenesis, i.e. when the first neuron is produced in the neocortex, and *t_F_* as the final time of neurogenesis, and consider the time interval *t ∈ (t_0_,t_F_).*

Cell death of post-mitotic neurons is not explicitly modelled, but will be accounted for when integrating experimental estimates of final neuronal output into the model. Similarly, the model can be refined to account for cell death of the progenitor cells. Gohlke et al. (2007) investigated the role of cell death in a computational model of neuronal acquisition.

A vast literature for cell cycle models is available (Weis et al. 2014), and different modelling approaches can be adopted for the description of the cell cycle at different scales of complexity. We are interested in the cell population dynamics, a higher scale level than the fine-grained study of progression through cell cycle phases and the molecular processes involved. Therefore, we simply account for the progenitor cell cycle length *T_C_*, defining the growth rate as: *ρ* = log 2/*T_C_*.

In the special case of constant *α* and *β* the *P* population either goes extinct, remains constant, or grows without bound, depending on the value of (1 –*α*–; 2*β*), as shown in supplemental Figure S1. Our time-dependent probabilities will model the observed patterns of division (Takahashi et al. 1996), with SymP the preferred mode of division at early neurogenesis, AsymN peaking at mid-neurogenesis and finally decreasing to leave SymN as the preferred mode of division at late neurogenesis. Therefore, we choose the probability functions to be piece-wise linear functions of time and define the following parameters: *α*_0_, the proportion of AsymN at onset of neurogenesis; *α*_S_, the proportion of AsymN at the time of strategy switch; *β_F_*, the proportion of SymN at the end of neurogenesis; and *t*_S_, the time of the switch to a strategy favouring SymN. These four parameters uniquely determine the shape of the probability functions represented in Figure 2, hence the trajectory in parameter space (Fig. 1). For an analytical description of the probability functions, see the supplemental information. We only consider biologically relevant parameter values, namely:

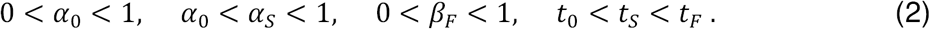

Within these constraints, any combination of the four parameter values will correspond to a trajectory in the strategy space.

**Figure 2.**
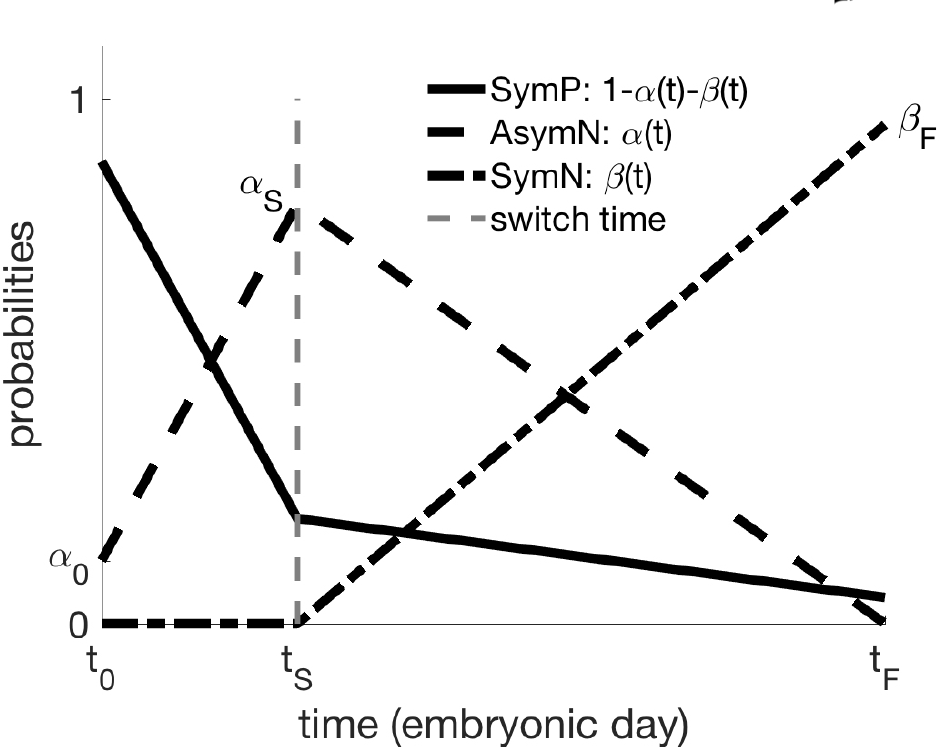
Time-dependent probability functions for the three types of division. A two-step linear strategy is hypothesized, with a shift from asymmetric to symmetric neurogenic division and concurrent reduction in self-amplifying divisions. Neurogenesis occurs in the time interval (*t_0_,t_F_).* Parameters *α_0_, α_S_, β_F_, t_S_* unequivocally determine the shape of the functions.

The model outcome, given a choice of parameter values, consists of the functions of time *P*(*t*) and *N*(*t*), representing the cell counts over the course of neurogenesis, i.e. for *t* ∈ (*t_0_,t_F_*).

We initially consider mouse embryonic cortical neurogenesis, occurring between E11 and E19 (E indicates embryonic day), and fixed cell cycle length, *T_C_* = 14.3 hours (see Table 1). However, in a second version of the model, we introduce an age-dependent cell cycle time, *T_C_*(*t*), to account for the experimentally observed modulation of the cell cycle length throughout the course of neurogenesis, mostly due to variations in the S phase (Dehay and Kennedy 2007).

## Results

Our study focuses on three species of interest: mouse, macaque monkey, and human. We intend to build a mathematical framework that can easily be extended to consider a larger number of species, with the aim of analysing and explaining cortical neurogenesis through the lens of evolution. Specifically, our study focuses on cortical neuronal number, rather than brain size. This allows us for a more straightforward comparison of proliferative and differentiative dynamics across species, without the need to introduce species-specific characterization of cell size and density. Although data on a large number of other mammalian species exist, we focus on these three species that have been chosen in previous studies to effectively illustrate neocortex development across distinct branches of the phylogenetic tree (Rakic 1995).

We initially search the (AsymN, SymN, SymP) space to find the strategy, or strategies, matching the observed patterns of cerebral cortical neuronal numbers in each species. Then, we use each species’ estimated strategy to make predictions about the size of the founder population, currently unavailable in the experimental literature. The model we propose can be used to compare different developmental strategies to determine the existence of evolutionary or developmental constraints. Finally, we can determine those parameters, to which the model outcome is most sensitive, suggesting the need for higher resolution data, and identify those parameters whose variation does not substantially affect model predictions.

### The strategy to build a mouse neocortex

The laminar position of a newly generated neuron is determined by the time of birth in an “inside-first, outside-last” fashion (McConnell and Kaznowski 1991), therefore we can assume that the radial expansion of the neocortex is proportional to the number of neurons. In mice, specifically, the total number of neurons per column is approximately shared between upper and deeper cortical layers, while upper layers are significantly expanded in primates (Markram et al. 2004; Vasistha et al. 2015; Charvet et al. 2015; Table 1).

Since we are interested in the ratio of population sizes at different times, we can rewrite equations (1) as:

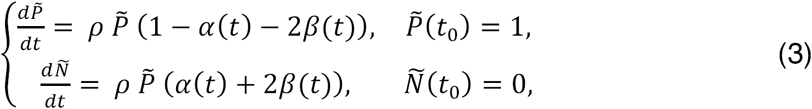

where 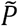 and *Ñ* are progenitor and neuron cell numbers, respectively, per initial progenitor *P_0_*, so that the problem is independent of the size of the founder population.

Therefore, we define the *mouse strategy* as the 4-tuple (*α_0_,α_S_,β_F_,t_S_*) that best approximates the following:

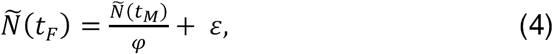

with minimum errors ε. Here, *φ* is the ratio of neurons in deeper layers (V and VI). *t_M_* is defined as the time when production of layer V is completed and production of layer IV starts. Hence, we look for the strategy that, to a first approximation, produces *φN*(*t_F_*) neurons by *t_M_*. Specifically, in the mouse cortical neurogenesis *φ* = 0.52 and *t_M_ =* E17 (see Table 1).

Given the small dimensionality of the problem, it is computationally feasible to perform a systematic and exhaustive search of the parameter space corresponding to all possible 4-tuples that satisfy equations (2). Doing so, we obtain the strategy:

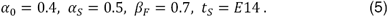

The trajectory corresponding to this mouse strategy is shown in Figures 3(A) and (C). Note that, given the linearity of the system, the strategy satisfying equation (4) for *ε* = 0 will still be valid for any value of *P_0_,* or for a different neurogenesis length (*t_F_ − t_0_),* provided that the time of switch *t_S_* is rescaled. Supplemental Figure S2 shows the result of the full search of the 4-dimensional parameter space.

**Figure 3.**
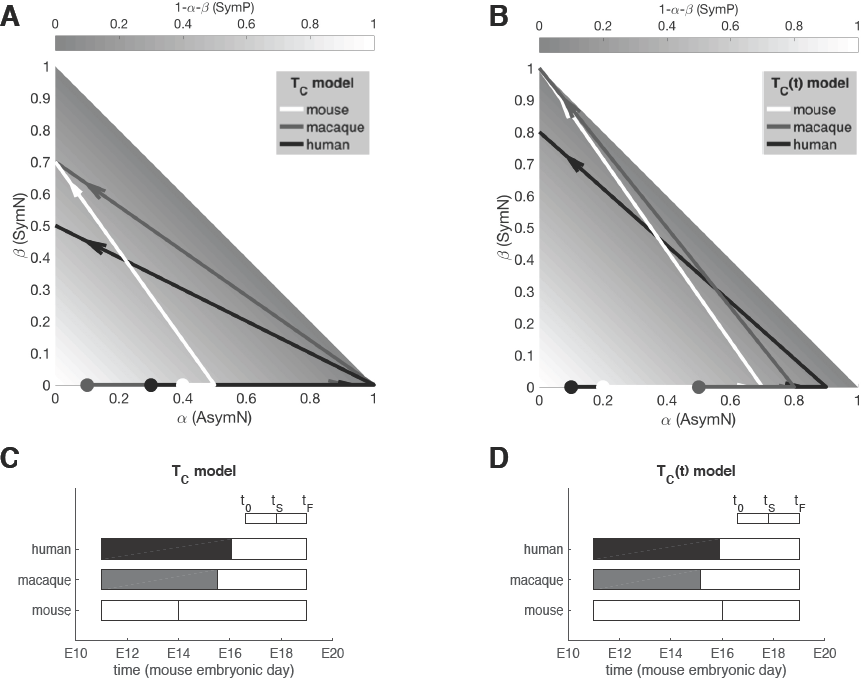
Representation in strategy space of the mouse, macaque, and human strategy for constant **(A)** and age-dependent **(B)** cell cycle length models. The lower panels, **(C)** and **(D)**, respectively, show the corresponding estimates for the time of switch. Macaque and human times are rescaled in the equivalent mouse time scale, to facilitate a meaningful comparison of the timing of events across species.

Our parameter sensitivity analysis (see supplemental information for details) reveals that the highest sensitivity of the output can be attributed to variations in the *t_S_* parameter (Fig. 4). This means that the variation in model output observed when varying the time of switch (*t_S_*) is more substantial when compared to the effect of varying the absolute prevalence of different types of division (*α_0_,α_S_,β*_F_). This sensitivity, as well as lack of experimental quantification, illustrates a demand for future experimental research into quantifying, or at least characterising, this time scale.

**Figure 4.**
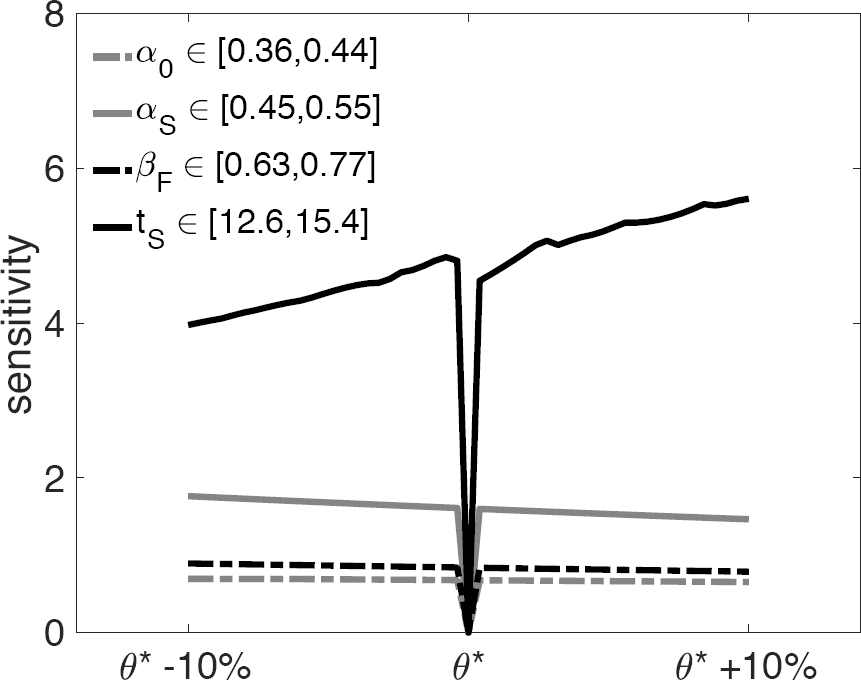
Local sensitivity analysis of strategy parameters around the mouse strategy (Eq. 5). Sensitivity is calculated according to the formula reported in the supplemental information. *θ\** indicates the reference value for the parameter whose sensitivity is being calculated. By definition sensitivity corresponding to *θ\** is zero.

### Species-specific strategies

We now move from the mouse, where neurogenesis has been extensively studied, to primates, where many key processes are relatively poorly characterized. Despite the lack of experimental quantification, we proceed with the best approximation found in the literature, and later test the robustness of our results against these simplifying assumptions and approximations. When evaluating the macaque strategy, we implement the method previously presented for the mouse, adjusting the timing of events (*t_0_,t_F_,t_M_*), the cell cycle length *T_C_,* and the ratio of deeper layer neurons *φ*. Finally, in order to evaluate the human strategy, values for parameters *T_C_* and φ, are approximated using the macaque parameters, as they cannot be found in the literature. A summary of model parameters adopted, and resulting strategies for each species, are reported in Table 1.

### Age-dependent cell cycle

A simplifying assumption in the model is that cell cycle length is kept constant during neurogenesis. However, species-specific age-dependencies could significantly affect the total number of divisions occurring during neurogenesis, and consequently the final neuronal output. For example, the linear increase of cell cycle length observed in mice determines a lower rate of division in late neurogenesis. In macaque, however, an initial increase in cell cycle length is followed by a decrease in mid-to-late neurogenesis (Kornack and Rakic 1998). This age-dependent modulation of frequency of division, combined with the temporal modulation of propensities of the modes of division, can substantially alter the dynamics and, hence, the outcome of neurogenesis. Our model (1) can be extended to account for this time dependency (i.e. *ρ* = *ρ*(*t*)) to give a more accurate representation of the system and describe what turn out to be non-intuitive implications, in terms of neuronal output.

For each species of interest we chose a linear description of *T_C_*(*t*), obtained by interpolation of the data from Kornack and Rakic (1998; see Fig. 5A top row, and supplemental information for the analytical definition). Repeating the search in the strategy space with the age-dependent model, calibrated to each species, we obtained a new set of strategies, reported in Table 1 and represented in Figures 3(B) and (D). Intuitively, the delay in the time of switch to neurogenic divisions predicted for the mouse strategy, increases the pool of progenitors to compensate for a decrease in the frequency of division at late mouse neurogenesis. Conversely, the non-monotonic cell cycle regulation throughout primate neurogenesis results in an earlier time of switch for both macaque and human.

**Figure 5.**
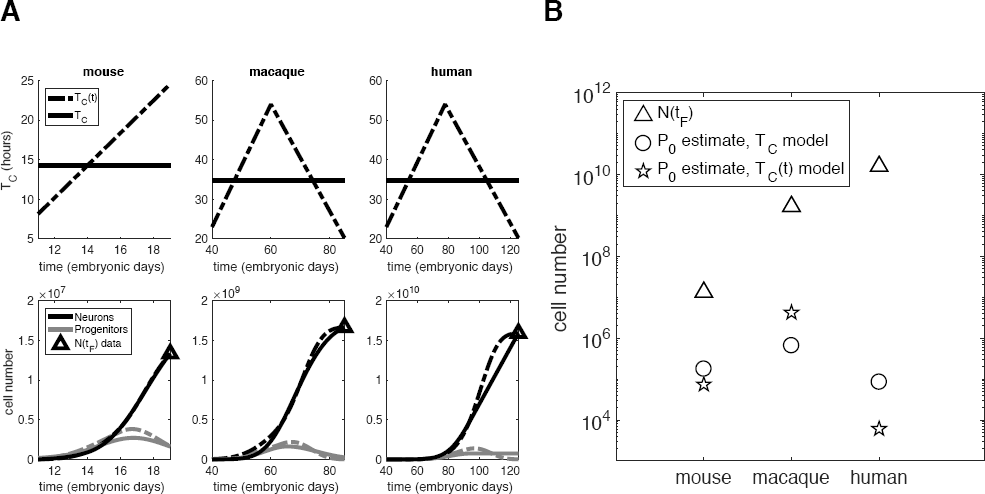
**(A)** Top row: constant (solid line) and age-dependent (dashed line) cell cycle models for mouse, macaque, and human. Bottom row: Solutions *N*(*t*) and *P*(*t*) for the two alternative cell cycle models parameterized on the three species. **(B)** Founder population estimates for mouse, macaque, and human using the corresponding strategy for constant (o) and age-dependent (☆) cell cycle length models.

### Estimate of founder population for mouse, macaque and human cortex

The number of cortical neurons present at the end of neurogenesis far exceeds the number in the adult neocortex, because of significant apoptosis of progenitors and newly born neurons (Kuan et al. 2000; Pompeiano et al. 2000). Additionally, in several instances, neurons migrate to the cortex after being generated elsewhere. Some examples are glutamatergic Cajal-Retzius and subplate neurons generated in the cortical hem or rostral medial telencephalic wall (Bielle et al. 2005; García-Moreno et al. 2007; Pedraza et al. 2014) or GABAergic interneurons generated in the medial, lateral and caudal ganglionic eminences (De Carlos et al., 1996; Lavdas et al., 1999; Anderson et al. 2001; Corbin et al. 2001; Tamamaki et al. 2003). Therefore, we estimate the neuronal output *N(t_F_*) adjusting the number of adult cortical neurons for 30% cell death and 25% of interneurons migrating to the cortex. Later we show that the results are robust of variations in these quantities.

Given *N*(*t_F_*), we intend to use our neurogenesis model to estimate a key quantity of interest that lacks experimental quantification: the founder population, *P*_0_. For each species of interest, we use Approximate Bayesian Computation (ABC, see supplemental information) to obtain an estimate of *P*_0_. This method gives the best fit to match the neuronal output *N*(*t_F_*), given the model of choice (constant, or age-dependent, cell cycle length), and the corresponding strategy. Figure 5(B) shows the estimates of *P_0_* for the three species and two models.

Both the constant and age-dependent cell cycle models predict that the exponential increase of cortical neurons from mouse to macaque is justified by a linear increase in the founder population (note the logarithmic scale on the *y* axis). This prediction agrees with the radial unit linear model (Rakic 2009), attributing an exponential expansion of the number of radial columns to the linear increase in the symmetrically dividing neural stem cell pool that, in our model, is represented by the founder population *P*_0_.

Our estimate for human *P*_0_, however, leads to a somewhat counterintuitive prediction. When moving from macaque to human, the founder population required to produce larger neuronal output is not increasing. Specifically, the founder population estimated for the human cortex is smaller than those values predicted for both mouse and macaque, despite producing a considerably larger number of neurons.

Solutions *N*(*t*) and *P*(*t*) for the parameterised models are shown in Figure 5(A), bottom row. It is worth noting that, despite the estimation being obtained by fitting one data point only, *N*(*t_F_*), for a given strategy, the time course of the model output does not vary significantly when comparing the two models (compare solid and dashed lines). Therefore, a more accurate representation of age-dependent cell cycle length does not change the time dynamics, but it does significantly affect quantitative estimations of the size of the founder population, especially for larger species.

To test the robustness of our result and challenge such a counterintuitive prediction, we introduce variations in the assumptions that could have skewed the results. Specifically, we repeat the estimations for varied amounts of interneuronal migration and post-neurogenesis cell death. Even an unrealistically large rate of post-neurogenesis cell death does not alter the prediction of lower human founder population (Suppl. Fig. S3A). Similarly, the prediction still holds even when increasing the rate of interneuronal migration to 50% (Suppl. Fig. S3B), although a more realistic 20% rate has been suggested (Petanjek et al. 2009). Finally, due to the lack of quantification of cell cycle length for human progenitors at various developmental stages, we used the experimental data for the macaque cell cycle length. This assumption, adopted in previous comparative studies (Rakic 1995), is valid only as a first approximation, since the two species belong to the same order. However, an underestimation of the cell cycle length in humans could cause an underestimation of the corresponding founder population. Additionally, a different frequency of cell divisions will alter the strategy adopted. Therefore, we re-evaluate the human strategy for longer cell cycles, up to twice the baseline value for macaque progenitor *T_C_* (Table 1), and subsequently repeat the corresponding estimations of the founder population (Suppl. Fig. S4A and B for constant and age-dependent cell cycle length models, respectively). For such large values of cell cycle amplification, the model prediction changes. The constant cell cycle model predicts that an increasingly large founder population is necessary to obtain increasingly larger neuronal outputs by the end of neurogenesis. The more accurate age-dependent cell cycle model, however, predicts that, even with a human cell cycle largely amplified with respect to the macaque, the founder population is approximately conserved.

### The neurogenesis simulator

We have developed a graphic user interface allowing the user to select and calibrate the neurogenesis model, observing how the temporal population dynamics vary when changing parameter values. The user can select and compare strategies of neocortical neurogenesis across different species. The neurogenesis simulator is available for download at www.dpag.ox.ac.uk/team/noemi-picco.

## Discussion

We propose a simple mathematical model of cortical neurogenesis that aims at describing the dynamics of cell proliferation and differentiation in various species.

Specifically, we can think of the cortex in each species arising as the result of a controlled balance of cell proliferation and differentiation, to obtain the developmental strategy. Such a strategy consists of the modulation in time of these processes during development (Fig. 1) so that, within the time allowed for cortical neurogenesis in each species, the required number of neurons is produced and cortical layers are formed.

In order to make meaningful comparisons between species, we defined a two-population model, describing the time dynamics of progenitors and neurons during cortical neurogenesis. The progenitor population is defined differently for any species of interest and its description can be refined to include different types of progenitor cells (e.g. neuroepithelial cells, radial glial, intermediate progenitors, etc.).

The main hypothesis of this model is that the three types of division (SymP, AsymN, SymN) are preferentially adopted at different stages of neurogenesis. Therefore, we imposed a two-step linear strategy with the prevalent mode of division shifting from self-replication at early neurogenesis to neurogenic differentiation at later times (Fig. 2). We propose two variants of the model, with constant, and age-dependent, cell cycle length, respectively.

The model framework allows the systematic exploration of the strategy space by simultaneously varying all parameter values within biologically relevant ranges to find the strategy (strategies) matching the observed outcomes (Suppl. Fig. S2). Thus we can obtain the developmental strategy for mouse, macaque, and human neurogenesis (Fig. 3).

Despite imposing the shape of probability functions, to describe a shift in prevalence of mode of division, we do not make assumptions about the timing of the switch from mostly self-replication to mostly neurogenic differentiation. The model prediction is that, to match the pattern of neuronal output observed, the time of switch must be around early/mid-neurogenesis in mouse, and around mid/late-neurogenesis in macaque and human (Fig. 3C). The introduction of a more accurate age-dependent cell cycle model results in a later time of switch in the mouse strategy, and an earlier time of switch in the macaque and human strategy (Fig. 3D).

We then used sensitivity analysis to determine which parameters have the highest influence on the model outcome. We found that the timing of switch of strategy from mostly self-replicating to mostly neurogenic divisions is more important, in terms of neuronal output, than the mere proportions of these types of division (Fig. 4). Although a number of cell-intrinsic factors have been found to regulate modes of division at the single-cell level, a full understanding of the mechanisms is lacking. For example, overexpression of p27^Kip1^, a cyclin-dependent kinase inhibitor, was found to substantially reduce neuronal output in supergranular layers (i.e. late-stage neuronal production; Tarui et al. 2005). Although this constitutes only circumstantial evidence, p27^Kip1^ is a potential candidate for the experimental validation of the effects of a delay in the time of switch. Similarly, *β*-catenin was found to be a key regulator of cell cycle re-entry of neural precursors, promoting self-replicating divisions and generating aberrant cortices in mutant mice (Chenn and Walsh 2002). Primary microcephaly is an example of a developmental disorder associated with a mistimed switch between proliferative and differentiative divisions. Specifically, all microcephaly genes identified are related to centrosomal abnormalities affecting mitosis (Gilmore and Walsh 2013).

Having characterised the species-specific strategies, we used the model to estimate another quantity of interest, currently unavailable in the literature, namely, the size of the founder population of progenitors. Our model makes a counterintuitive prediction: the number of founder progenitors needed to produce the neurons present in the human cortex at the end of neurogenesis is lower than the corresponding estimate for macaque, despite the two-fold increase in neuronal output (Fig. 5). This prediction holds even for large variations in key parameters, such as rates of post-neurogenesis cell death and interneuronal migration (Suppl. Fig. S3).

Finally, we investigate the effects of variations in cell cycle length, assumed to be approximately similar in all primates, given the lack of data. We allowed some variation from the baseline value (macaque) and observed that the model prediction breaks down for large, unrealistic, values of human progenitor cell cycle length. This is an example of how, by integrating theoretical and experimental modelling, we can determine how other observed phenomena affect our hypotheses, predictions and understanding of the process that is being investigated. This approach is particularly relevant in resolving questions arising from circumstantial evidence. Indeed, the theoretical model can suggest whether a given mechanism could lead to significantly different predictions, suggesting that further experimental investigation is necessary, or otherwise futile for the problem being investigated.

Mora-Bermúdez et al. (2016) estimate an approximate 1.3x amplification in progenitor cell cycle length in human organoids, with respect to macaque. To the best of our knowledge, however, a quantification of the cell cycle length of human progenitors is not available from the current literature in vivo. Therefore, at this stage, we can only speculate on the implications of this counterintuitive prediction. However, our study suggests that further experimental studies should focus on the quantification of cell cycle duration, as our model shows that very different predictions can follow. Conversely, variations in other properties of the system, such as post-neurogenesis neuronal cell death and interneuronal migration, do not affect the qualitative predictions of the model.

There is strong evidence of a link between cell cycle lengthening (specifically, of the G1 phase) and the progression to a higher propensity to differentiative divisions (Dehay and Kennedy 2007). However, the relationship of causality between these two factors has yet to be established. By proposing two versions of the cell cycle model, we offer an alternative interpretation of the developmental program, with no direct relationship between cell cycle length and preferred modes of division. Our constant, and age-dependent, cell cycle models lead to qualitatively similar predictions regarding the scaling of the founder population, but very different quantitative predictions for a given species (Fig. 5B). Experimental quantification of *P*_0_ would give the added benefit of validating either one of the two models, determining the need (or not) of age-dependent cell cycle to quantitatively recapitulate the population dynamics.

### Conclusions

With this study we introduced a minimal theoretical framework which, informed by experimental observations, offers a mechanistic explanation of the development and evolution of the mammalian neocortex. This model allows a comparative study of mammalian species, and future modelling of human pathologies, such as microcephaly. In building our first minimal model we left out a number of properties of the system and mechanisms that could be integrated as they become relevant and necessary to answer specific biological questions, and as data becomes available. For example, radial migration is only modelled implicitly, by assuming that positioning in the radial direction is related to time of birth (Rakic 1974; McConnell et al. 1991) fundamental to establishing a relation between the timing of neurogenesis and the distribution of neurons between upper and deeper cortical layers (Eq. 4). Additionally, our model captures the dynamics of neuronal acquisition averaged over the entire neocortex, neglecting variations in cortical areas. Early regionalization determined by variations in cell-cycle kinetics is crucial (Polleux et al. 1997; Dehay and Kennedy 2007). However, such differences are relatively minor compared to the cross-species variation that we intended to address with this work. Furthermore, our model explains inter-species variation in neuronal output regardless of cortical surface and folding. To explain these differences (i.e. gyrencephalic vs lissencephalic) we would have to account for density, cell size, tangential migration, etc., all of which are species-specific quantities and properties. Specifically, it was shown that cortical folding approximately scales with surface area, and thickness, and not with neuronal numbers (Mota and Herculano-Houzel 2015). Additionally, through integration of imaging and mathematical modelling it was demonstrated that variations in cortical thickness within the cortex are caused by the physical forces emerging during the folding process (Holland et al. 2018). However, the problem of neocortex folding is a separate avenue of research, decoupled from the mechanisms that we want to investigate here.

The aim of this work was to go beyond scaling rules, to identify the minimum set of players, properties and interactions sufficient to explain cross-species variation in cortical neurogenesis in terms of mechanisms.

Specifically, we can exploit the predictive and analytical power of mathematical modelling to discriminate between the many factors contributing to the formation of the cortex. Additionally, sensitivity analysis allows us to pinpoint those mechanisms where higher resolution data would be beneficial to improve our understanding of cortical neurogenesis. The usefulness of mathematical modelling is two-fold. It can be used to guide us on experiments that are necessary to quantify the key parameters that need to be systematically compared across species. To this end we introduced the *neurogenesis simulator,* a tool allowing the user to observe changes in the model outcome as model parameters and assumptions are varied. Mathematical modelling can also be helpful in formally proving (or falsifying) empirical hypotheses generated from experimental observations, such as the radial unit hypothesis formulated many decades ago by Rakic (1988). We found that the timing of shift from self-replicating to neurogenic divisions is key to obtaining the correct neuronal numbers by the end of neurogenesis. The lack of experimental quantification of a second key quantity, cell cycle duration, leaves an open question. Namely, whether the predicted reduction in founder population size of human versus macaque is real. If experimentally validated, this prediction would have a key implication in evolutionary terms. That is, species with increasingly larger numbers of cortical neurons could likely have adopted the same *developmental strategy* adjusting the duration of neurogenesis, thus requiring an approximately conserved size of founder population. Indeed, a founder population size that scales with the neuronal output would be prohibitively large and inefficient, compared to running the same developmental program (strategy) over a stretched window of time.

We envisage the use of our neurogenesis model to map evolutionary trajectories that describe neurogenesis in different species, as well as deviations in such trajectories corresponding to brain diseases characterised by a developmental program gone awry.

## Acknowledgements

This work was supported by a St John’s College Research Centre Grant to TW, PKM and ZM in collaboration with FG-M. The work in the laboratory of ZM was supported by grants from MRC, Wellcome Trust, Royal Society and St John’s College Research Centre. PKM and TEW would like to thank the Mathematical Biosciences Institute (MBI) at Ohio State University, for partially supporting this research. MBI receives its funding through the National Science Foundation (grant number DMS1440386). FG-M holds an IKERBASQUE Research Fellowship. The authors gratefully acknowledge useful discussions with members of the Brain Research And Inquiry Network (BRAIN) at St John’s College Research Centre. We also thank Dr. Navneet Vasistha, Dr. Patricia Garcez, Prof. Bruno Mota, Prof. Alain Goriely for discussion, critical reading and feedback on earlier versions of this manuscript. We would like to thank the three reviewers for their valuable feedback and constructive comments.

